# FGF13 is not secreted from neurons

**DOI:** 10.1101/2025.11.12.688075

**Authors:** Mattia Malvezzi, Haiying Zhang, Patrick Towers, David C. Lyden, Steven O. Marx, Geoffrey S. Pitt

## Abstract

FGF13, a noncanonical fibroblast growth factor (FGF) and member of the fibroblast growth factor homologous factor (FHF) subset, lacks a signal sequence and was previously reported to remain intracellular, where it regulates voltage-gated sodium channels (VGSCs) at least in part through direct interaction with the cytoplasmic C-terminus of VGSCs. Recent reports suggest that FGF13 is secreted and regulates neuronal VGSCs through interactions with extracellular domains of integral plasma membrane proteins, yet supportive data are limited. Using rigorous positive and negative controls, we show that transfected FGF13 is not secreted from cultured cells in a heterologous expression system nor is endogenous FGF13 secreted from cultured neurons. Further, employing multiple unbiased screens including proximity protein proteomics, our results suggest FGF13 remains within membranes and is unavailable to interact directly with extracellular protein domains.

## INTRODUCTION

For nearly 30 years since they were first discovered, fibroblast growth factor homologous factors (FHFs) have been described as members of the FGF subfamily that differs from canonical FGFs in two key aspects: they lack a signal peptide and therefore are not secreted; they do not bind and activate FGF receptors at physiological concentrations (1). Instead, they have been shown to act intracellularly. There are four FHFs, FGF11-14 (FHF3, 1, 2 and 4, respectively), and while their best documented role is the regulation of voltage-gated sodium channel (VGSC) activity (2–11), achieved via direct binding to the VGSC C-terminal domain (8, 12–15), recent data support VGSC independent roles, such as microtubule stabilization (16) and synaptic transmission regulation (17). Furthermore, FHFs, specifically FGF13, are overexpressed in many tumors (18–20), although their role remains unclear.

FHF genes generate multiple isoforms via alternative splicing (21). Among the FHFs, the brain expresses predominantly FGF13-S (aka FGF13-A or FHF2A) and FGF13-VY, with FGF13-S expressed mainly in excitatory neurons (22, 23). FGF13-S is highly concentrated at the neuron axon initial segment (AIS), where it regulates VGSC current density (10).

Several recent studies challenged the paradigm that FHFs are not secreted and do not activate FGF receptors (24–27). Specifically, FGF13-S was suggested to be secreted from neurons, causing inhibition of VGSCs and a reduction in the intrinsic excitability at the AIS via binding to the extracellular receptor LRRC37B, with implications for epilepsy and neurodevelopmental disorders (27). FGF13-S secretion, however, was analyzed only in a heterologous system and without the benefit of a complement of positive and negative controls. Whether FGF13-S is secreted from neurons and has a consequent extracellular physiological role is not known, thus clouding an understanding of its physiological functions.

Here we show that FGF13-S is not secreted from HEK293 cells, even in the presence of stress, nor from isolated mouse hippocampal neurons, while known secreted proteins were readily found in the extracellular space. Any FGF13-S detected within the extracellular medium of cultured HEK293 cells was restricted to large extracellular vesicles (LEVs) and, to a lesser extent, to small extracellular vesicles (SEVs), which include exosomes. In an orthogonal analysis, we performed *in vivo* proximity labeling proteomics in the mouse brain and found that FGF13-S near neighbors are cytoplasmic, non-secreted proteins. These results provide clarification that FHFs remain cytoplasmic and do not function as secreted proteins.

## RESULTS

Assays for determining whether a transfected protein is secreted from cultured cells and available to bind to extracellular receptors generally rely on detecting the protein in the extracellular medium of the transfected cells after clearing the medium of dead or dying cells and of extracellular vesicles, in which the queried protein may be present yet unavailable to interact directly with extracellular receptors. To test if FGF13-S is secreted from HEK293 cells (in which FGF13-S is not endogenously expressed), we transiently transfected FGF13-S and then employed a multi-step differential centrifugation protocol designed to minimize contamination from FGF13-S released from dead or dying cells or FGF13-S within extracellular vesicles. We also transfected hemagglutinin (HA)-tagged adipsin, an adipokine secreted from adipose tissue (28), as a positive control for secretion. Cells were cultured for 24 hours after transfection with either FGF13-S or adipsin and then incubated for 24 hours in fresh, serum-free medium. Serum was removed for two reasons: i) to allow us to concentrate the cleared extracellular medium and load comparable amounts of protein among the different fractions, which is not possible when serum is present because of the large amount of albumin which creates artifacts when running the samples on a polyacrylamide gel, even when the samples are not concentrated (Suppl. Fig. 1A); ii) to reproduce the experimental conditions in previous studies in which FGF13-S was reportedly detected in the serum-deprived extracellular medium of cultured cells (25). Extracellular medium was collected and subjected to sequential centrifugations. A first, low-speed (4000 × g) centrifugation cleared the medium of any cell debris. The pellet was discarded, and the cleared medium was then centrifuged at 12,000 × g to precipitate large extracellular vesicles (LEVs) and then at 100,000 × g to collect small extracellular vesicles (SEVs), including exosomes. LEVs and SEVs were resuspended, lysed, and loaded on gels for western blot analysis together with the concentrated (30×) cleared medium, in which true secreted proteins are expected. Indeed, abundant adipsin signal was detected in the cleared medium, but FGF13-S was not (Fig. 1A). It is worth noting that we not only detected adipsin in the extracellular medium, but that it has a higher molecular weight (∼40-50 kDa) than the adipsin detected in the cell lysate, reflecting the processing (glycosylation) of the intracellular, pre-secreted 28 kDa protein (28), (the apparent molecular weight of the protein is larger than native adipsin due to the presence of the HA tag), providing validation for our medium-clearing protocol. The broad band spanning ∼10 kDa observed in the cleared medium likely reflects variable glycosylation states (28). In addition to detecting FGF13-S in the cell lysate, we detected FGF13-S in LEVs and, to a lesser extent, in SEVs while adipsin was barely detected in either fraction. We further confirmed the identity of the extracellular vesicle (EV) fractions with validated markers. Specifically, Alix and CD9, two well-established exosomal markers (29), were found in the SEVs/exosome fraction but not in LEVs nor in the cleared medium (Fig.1A). Calnexin, an endoplasmic reticulum membrane protein, is observed in the cell lysate but not found in any fraction (Fig. 1A), confirming the successful removal of cell debris or cytoplasmic-only proteins from the EVs and medium. Thus, our clearing protocol enriches for LEVs, SEVs, and cleared extracellular medium while eliminating cytoplasmic debris from dead or dying cells. Taken together, these results show that FGF13-S is not secreted from HEK 293 cells but it is found in EV vesicles, possibly explaining its detection in previous secretion studies that did not utilize sequential centrifugation to separate EVs from extracellular material (25, 27).

**Figure 1.**
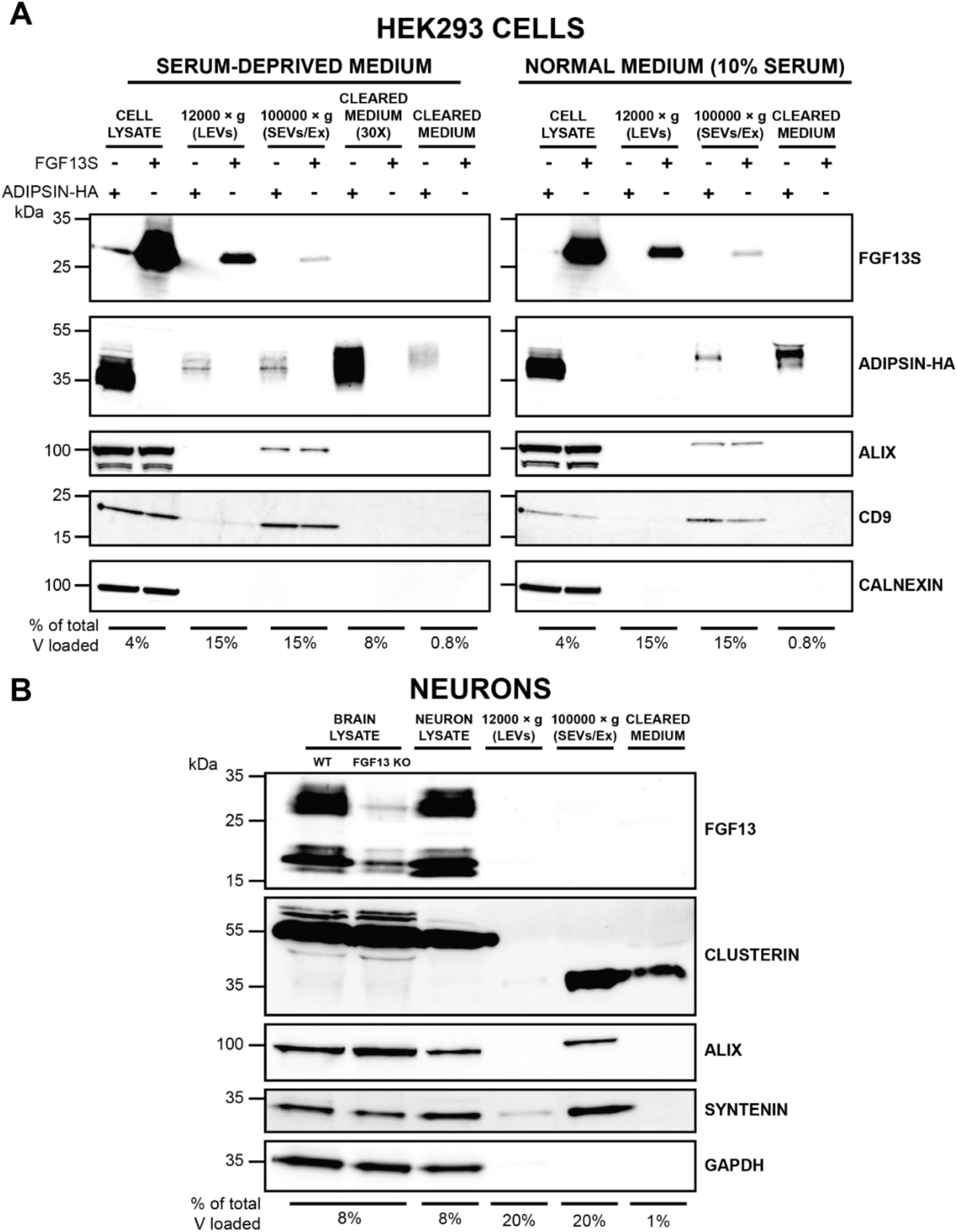
FGF13-S is not detected in cleared extracellular medium of HEK293 cells or culture mouse neurons. **A.** Western blot of cell lysates, extracellular vesicles, and cleared medium derived from HEK293 transiently transfected with FGF13-S or the secreted protein hemagglutinin-tagged adipsin (Adipsin-HA) and subjected to multi-step centrifugation protocol in serum-deprived conditions (left) or in normal culture conditions (10% serum, right). Alix (96 kDa) and CD9 (24 kDa) are used as SEVs/exosome markers; calnexin (90 kDa) as an intracellular marker. Secreted adipsin-HA has higher molecular weight as a consequence of post-translational modifications (28). Cell lysate (8 µg) was loaded. **B.** Western blot of cell lysates, extracellular vesicles, and cleared medium from cultured mouse hippocampal neurons after 12 DIV in culture. Whole brain lysates from wild-type and FGF13 brain KO were used to validate the detection of FGF13 isoforms. Clusterin is used as a positive control for secretion in neurons (non-secreted clusterin precursor: 55 kDa; cleaved, secreted clusterin: 35 kDa). Alix (96 kDa) and syntenin (34 kDa) are used as SEVs/exosome markers; GAPDH (36 kDa) as an intracellular marker. Fifteen µg of brain or neuronal lysate were loaded. % of total V loaded indicates the fraction of the sample loaded on each gel relative to the total volume.

As these experiments were performed under conditions of serum deprivation for the technical reasons noted above, the cells were under stress conditions. While this strategy was integral to previous studies in which FGF13-S was detected in the extracellular medium (25), we asked whether stress was required for HEK293 cells to release FGF13-S in EVs by also performing the sequential centrifugation protocol on cells for which the 10% serum was not removed. While the amount of cleared medium able to be loaded on the gel was 10-fold less due to the presence of serum components, primarily large amounts of albumin, that interfere with subsequent western blotting, we still clearly detected adipsin in the extracellular medium (Fig. 1A, right). For direct comparison between the serum-free and the serum-maintained conditions, we loaded the same reduced fraction (0.8%) of extracellular material and observed secreted adipsin for both conditions (Fig. 1A, left). Importantly, detection of FGF13-S in both LEVs and SEVs was unaffected by the presence or absence of serum, as was the detection of the positive and negative markers for EVs (Fig. 1A). Taken together, these results show that FGF13-S can be detected in EVs of HEK239 cells independent of stress; and that FGF13 is not detected in the cleared extracellular medium. Thus, we conclude that FGF13-S is not secreted from HEK293 cells.

The proposed physiological role for secreted FGF13-S is to regulate VGSCs in the brain so we specifically assessed whether FGF13 is secreted from neurons. Hippocampal neurons from WT mouse brain (P0) were cultured for 12 days (12 DIV), allowing neuronal maturation and testing for secretion. After 12 DIV, the medium was collected, cell debris was removed via centrifugation at 4000 × g and the extracellular medium was cleared via the same sequential spin protocol employed for HEK293 cells to separate LEVs (12,000 × g) and SEVs (100,000 × g). Cleared extracellular medium was not concentrated due to the high amounts of supplements in the medium (albumin and hormones) necessary to culture neurons, but which generate gel artifacts (Suppl. Fig. 1B). Cell lysates and the fractions obtained from the sequential spins were analyzed by western blots. Whole brain lysates from WT or neuronal-specific *Fgf13* knockout (*Nes-cre;Fgf13^fl/Y^*) mice provided validation of the antibody for detection of FGF13 brain isoforms, including FGF13-S. GAPDH, a cytoplasmic protein, was not detected in the extracellular medium nor in EVs (Fig. 1B), confirming successful removal of cell debris. Alix and syntenin, two exosomal markers (29), were readily detected in the SEVs fraction but not in the cleared medium, showing successful separation of EVs. Similar to the HEK293 cell experiments, the lack of detection of the cytoplasmic marker and the exosomal markers in the cleared medium demonstrates that the cleared medium is free from cellular debris and EVs. In contrast, clusterin, a disulfide-linked heterodimeric protein conventionally secreted from neurons (30, 31) and that undergoes cleavage and glycosylation of its 55 kDa precursor (32), was detected in the cleared extracellular medium (Fig. 1B). Consistent with previous reports (33, 34) we also detected clusterin in the SEV fraction. We additionally established that clusterin was secreted and accumulated in the extracellular medium because removing the medium after 11 DIV and replacing it with fresh medium for 1 additional day before collection yielded too little clusterin accumulation for detection (Suppl. Fig. 2). Under these conditions, we did not detect FGF13-S (nor any other known FGF13 splice variant) in the medium even though multiple FGF13 isoforms are present in neuronal lysates (Fig. 1B). In contrast to the HEK293 experiments, we did not detect any FGF13 in the EV fractions, suggesting that its presence in EVs might be cell type/tissue specific or dependent on stimuli such as stress. Together, these results do not support the hypothesis FGF13-S is secreted from neurons or that FGF13-S is released into the extracellular space from exosomes.

To provide an alternative means to assess if FGF13-S is secreted from neurons, we performed *in vivo* proximity labeling proteomic profiling in mouse brain. We reasoned that if FGF13-S is secreted, either through the conventional pathway or via a non-conventional pathway, it should be in the vicinity of proteins involved in these processes or other secreted proteins, and that at least a subset of those proteins would be detected by proximity labeling. Indeed, *in vivo* proximity labeling proteomic profiling has been used successfully to characterize secreted or extracellular proteins in various tissues and organs, including the brain (35, 36). We therefore generated transgenic mice with a cDNA encoding the biotin ligase TurboID (37) and a hemagglutinin (HA) tag fused to the N-terminus of FGF13-S under the control of a tetracycline-inducible promoter. We note that placing TurboID at the N-terminus could prevent FGF13-S secretion by blocking a signal sequence, yet a distinguishing feature of FGF13-S (like other members of the FHF subfamily) is the absence of a signal sequence (38). These mice were crossed with mice carrying a brain-specific (neurofilament heavy polypeptide, NEFH, promoter), tetracycline-controlled transactivator protein (tTA) transgene to achieve expression in the brain. We first validated the TurboID-FGF13-S expression in isolated hippocampal neurons by staining for HA (Suppl. Fig. 3). The transgene expression pattern recapitulates the somatic and axon initial segment enrichment of endogenous FGF13-S (23), albeit with an overall stronger signal due to overexpression. After *in vivo* biotin injections (or vehicle control) to promote TurboID-FGF13-S labeling of near neighbors, we captured biotinylated proteins with streptavidin and quantified them by mass spectrometry. We detected robust protein labeling in brains from mice injected with biotin compared to DMSO-injected mice (Suppl. Fig. 4A). Quantitative mass spectrometry analysis detected 1568 proteins (Supplemental Dataset, Suppl. Fig 4B), 457 of which were significantly enriched in biotin-injected samples (p<0.05 and Log_2_(Biotin/DMSO)>1) (Fig. 2A). Among the most enriched proteins was Na_V_1.6 (encoded by *Scn8a*), consistent with FGF13-S’s known concentration in the AIS (10) and interaction with VGSC cytoplasmic C-termini (13, 14). Other AIS enriched proteins, such as ankyrin G (*Ank3*) and β4 spectrin (*Sptbn4*), the potassium channels KCNQ2 (*Kcnq2*), K_V_2.1 (*Kcnb1*), and K_V_2.2 (*Kcnb2*), MAP7D2 (*Map7d2*), and Mical3 (*Mical3*) (39, 40) were prominently enriched in the dataset (Fig. 3A). Other prominent AIS proteins such as Gephyrin (*Gphn*), Contactin (*Cntn1*), Synaptopodin (*Synpo*), α2 Spectrin (*Sptan1*) and Prickle 2 were also identified, although slightly above the p<0.05 threshold. Sodium channel regulatory subunit β1 (*Scn1b*), a VGSC beta subunit to which secreted FGF13-S is proposed to cluster around via LRRC37B binding (27), was not detected.

**Figure 2.**
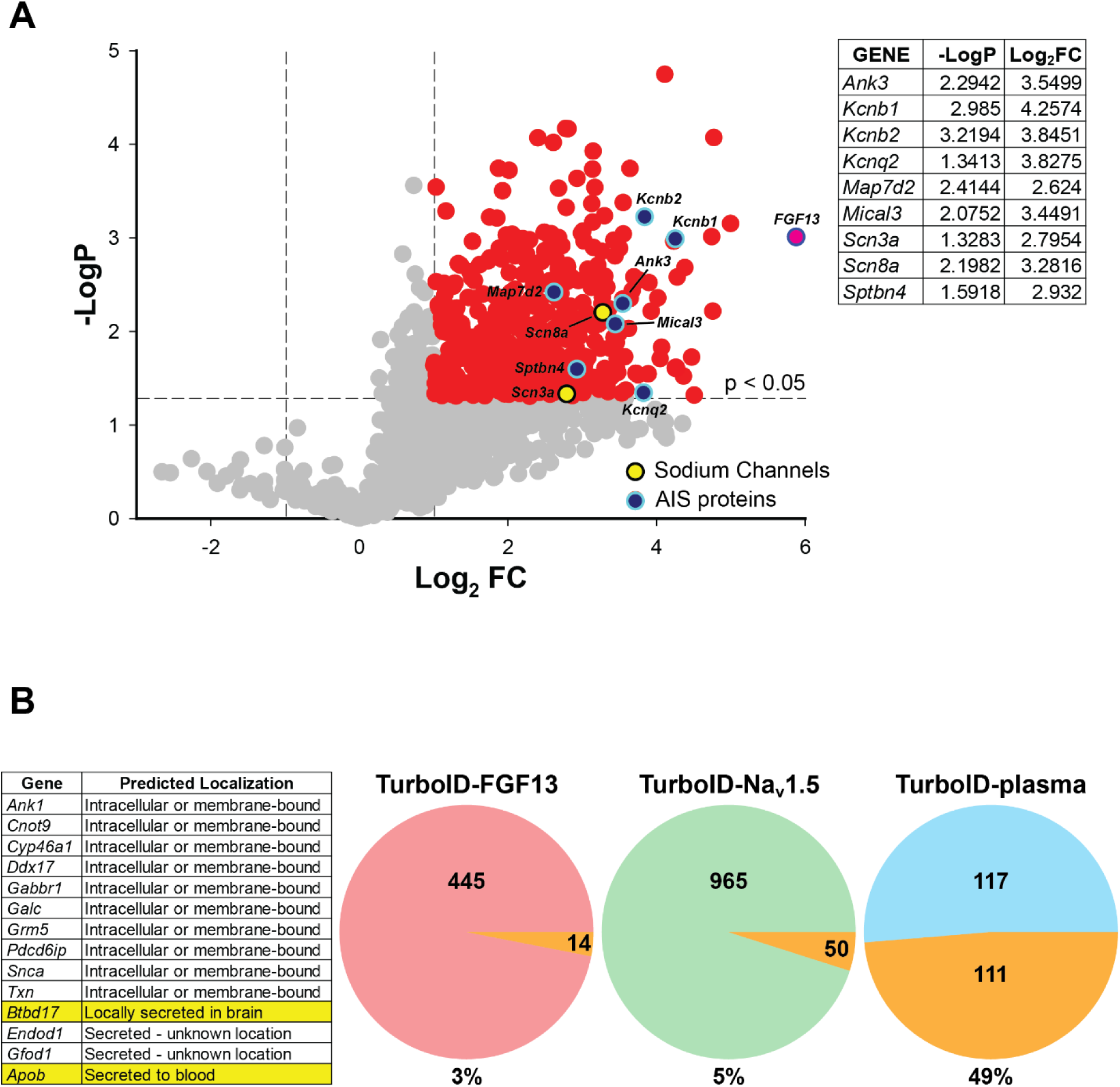
*in vivo* brain TurboID proximity proteomics shows FGF13-S is not in the vicinity of secreted proteins, nor proteins involved in secretion processes. **A.** Volcano plot of the fold-change of proteins enriched in biotin-injected animals over control. Red: highly enriched proteins, with a fold-change >2. Pink: FGF13. Yellow: sodium channels. Blue: axon initial segment proteins. Statistics for these proteins are shown in the table. **B.** Pie charts showing the overlap between the human secretome database of secreted protein (41) and our TurboID-FGF13-S (red) dataset, the TurboID-Na_v_1.5 dataset (42), (green) and a dataset of secreted protein detected by TurboID proximity proteomics (cyan, (36)) datasets. The full list of overlapping proteins found in our study, with their predicted localization, is shown in the left table.

**Figure 3.**
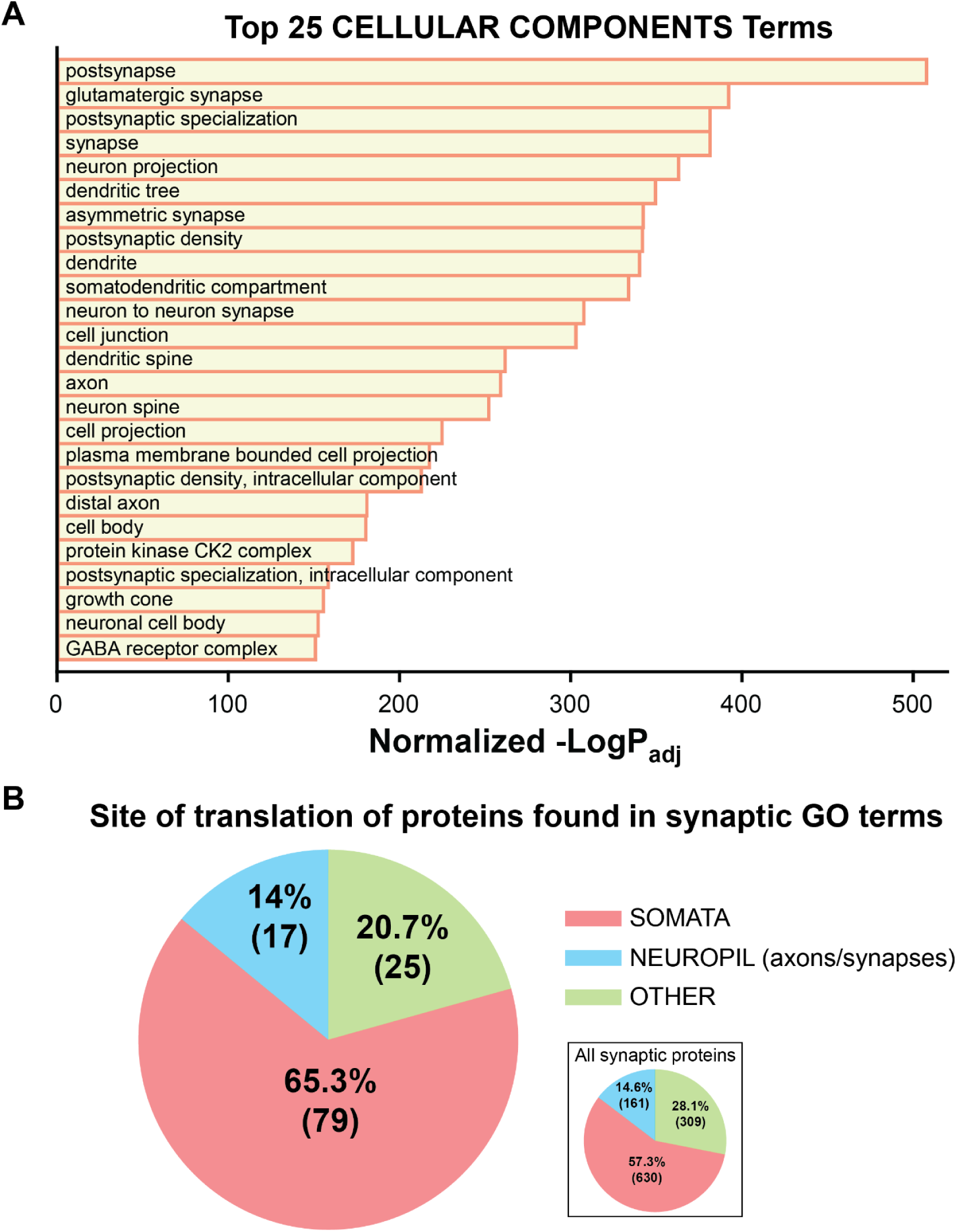
FGF13-S is in proximity of soma-translated synaptic proteins. **A.** Top 25 cellular component terms from gene set enrichment analysis showing a high representation of synaptic and distal axonal terms. Statistics are provided in the *Methods* section. **B.** Pie chart showing the majority of the proteins found in the cellular component terms shown in **A** are translated in the soma and not locally at the synapses. Sites of translation were obtained from Glock *et al.* (51). Thirty-two proteins were not present in the translation database; therefore 131 proteins were annotated. Inset: site of translation of all synaptic proteins, defined as the 2025 proteins comprising the *mus musculus* GO:0045202 term “synapse”. 925 proteins are not present in the translation database; therefore 1100 proteins were annotated.

With this validation of the dataset, we then asked whether any candidate FGF13-S near-neighbors are secreted proteins, which could suggest that FGF13 is similarly processed for secretion, despite its lack of a signal peptide. We exploited a recent study by Uhlen *et al*. colleagues that identified the human secretome (41), compiling a list of all proteins that are secreted, either extracellularly or intracellularly. The latter represents proteins going through the ER-Golgi canonical pathway but retained in intracellular organelles or vesicles. Of the 457 proteins significantly enriched in the FGF13-S TurboID dataset, 14 (3%) are present in the secretome database, and only 2 (APOB and BTBD17) are designated as extracellularly secreted (we excluded proteins that are predicted to be secreted intracellularly or that have an unknown location) (Fig. 2B). To assess the relevance of identifying 3% of proteins in the secretome database, we performed a similar analysis on a TurboID dataset for a known non-secreted protein, the cardiac sodium channel Na_V_1.5, obtained in cardiomyocytes (42). Among the 1015 proteins identified as Na_V_1.5 near-neighbors, 50 (4.9%) were also present in the secretome database, and 12 were annotated as secreted extracellularly. Thus, the 3% of proteins in the FGF13 TurboID dataset identified in the secretome appears to be within the range expected for non-specific detection of secreted proteins. In contrast, the overlap between a different TurboID dataset designed to detect proteins secreted from various cell types (36) and the human secretome dataset was 48.7% (Fig. 2B).

Additionally, we performed gene set enrichment analysis (GSEA) on the candidate FGF13 near neighbors and found that gene ontology (GO) terms with “secretion” were not among the top 100 biological processes identified (Supplemental Dataset). Further, for those GO terms with “secretion” in their identifiers, there were 52 members (proteins) detected by TurboID. Analysis of these 52 proteins showed that all were annotated as cytoplasmic or nuclear: none as secreted proteins (Suppl. Fig. 5). Further, we note that two key proteins necessary for the unconventional secretion of FGF2, TEC kinase and ATP1A1, are not among the significantly enriched FGF13-S neighbors. Since FGF13-S (like all FHFs) lacks a signal sequence, and secretion of the homologous FGF12 was proposed to utilize this unconventional pathway, the absence of TEC kinase and ATP1A1 among the candidate FGF13-S near neighbors suggests that FGF13-S is unlikely to be secreted in a FGF2-like fashion. Rather, the 52 candidate FGF13-S near neighbors found in GO terms with “secretion” in their identifiers are most likely cytoplasmic proteins that participate in secretory or other types of vesicle formation. Indeed, analysis of GO terms for Biological Processes or Cellular Components identifies multiple terms containing “vesicle” among the top hits (Suppl. Fig. 6). In a recent study of the related FGF13-VY in cardiomyocytes, we found that FGF13-VY regulated trafficking of Cx43 (*Gja1*) containing vesicles via affecting microtubule stability (42). Thus, FGF13-S may perform similar vesicle trafficking roles in neurons, and the identification of FGF13 in LEV and SEV fractionation experiments (Fig. 1) is consistent with a role of FGF13 in regulating vesicle formation and/or vesicle trafficking.

## DISCUSSION

Although FHFs are members of the fibroblast growth factor superfamily, for which almost all members are proteins secreted through the conventional pathway and activate growth factor receptors through interaction with the receptors’ extracellular ligand binding domains (43), the initial report identifying FHFs noted the absence of signal sequences and explicitly reported that FGF12 (FHF-1) was not secreted into the extracellular space (38). Subsequent studies identified various cytoplasmic FHF binding partners, including the cytoplasmic C-terminal domain of various VGSCs (2, 3, 12), the MAP kinase scaffold IB2 (44), microtubules (16), junctophilin 2 (45), and cavins (46). Lately, several studies have challenged the hypothesized restriction of FHFs to the cytoplasm, suggesting that FHFs (specifically, FGF12-A and FGF13-S) can be secreted extracellularly, where they locally modulate biological processes by binding to the extracellular domains of integral membrane proteins (24, 25, 27). Although FGF13-S has been extensively characterized as a regulator of VGSCs via binding to the channels’ intracellular C-terminal domain (47) and more recently by affecting local membrane cholesterol content (48), Libe-Philippot et al. reported that FGF13-S was secreted from neurons and bound to the extracellular domain of the human-specific protein LRR37B, resulting in modulation of Na_V_1.6 sodium channels at the AIS (27). In that study, however, results supporting FGF13 as an extracellular protein relied solely on the detection of FGF13-S in the extracellular medium after expression of FGF13-S in HEK293 cells, albeit without employing specific steps to eliminate FGF13-S released from dead/dying cells or FGF13-S within extracellular vesicles. Further, in the absence of comparison to a validated secreted protein, the relative efficiency of FGF13-S secretion could not be assessed. Whether FGF13-S is secreted from neurons was not tested.

Here, we tested if FGF13-S can be secreted from HEK293 cells using an analogous experimental setup while employing validated positive and negative controls and adding differential centrifugation steps to reduce contamination from apoptotic cells and separate EVs. We were unable to detect secretion despite finding abundantly expressed adipsin, a conventionally secreted protein. Rather, differential centrifugation showed some FGF13-S resides within LEVs and SEVs. We also tested if FGF13-S is secreted from cultured hippocampal neurons. While we detected robust secretion of clusterin, a positive control for an endogenously secreted protein, we observed no endogenous FGF13 isoform secretion, including FGF13-S. Further, by analyzing a database of proteins detected as FGF13-S near neighbors via TurboID, we obtained no evidence of FGF13-S secretion. Thus, with positive and negative controls, our results suggest that FGF13-S is not available to bind LRRC37B and thereby regulate Na_V_1.6 from the extracellular side of the membrane, as proposed (27). We conclude that FGF13-S remains in the cytoplasm. While the Libe-Philippot et al. study showed regulation of neuronal excitability after application of purified FGF13-S (or the N-terminal peptide encoded by the S exon), the biological relevance of those results is unclear if FGF13-S is not secreted. Additionally, those experiments did not employ negative controls (e.g., a scrambled peptide). Whereas here we specifically focused on the FGF13-S isoform, our use of a pan-FGF13 antibody in the cultured hippocampal neuron experiments suggests that none of the FGF13 splice variants in the mouse brain are secreted.

Beyond its well-characterized regulatory effects on VGSCs, recent work from others and us showed that FGF13 affects K^+^ channels (23, 49), controls local membrane cholesterol (42), stabilizes microtubules (16, 42, 50), and traffics ion channels and gap junctions to the plasma membrane (42, 48). Our results add to previous reports showing that FGF13 controls ion channel trafficking and suggest a potentially broader role in trafficking. Specifically, gene set enrichment analysis reveals that FGF13-S appears to be a near neighbor of an unexpectedly large quantity of synaptic proteins. “Synapse” is the top among the Cellular Components terms, and it comprises 153 (33.5%) of all the proteins identified. Several synapse-related terms, both pre- and post-synaptic, are among the top 25 terms (Fig. 3A). Yet previous studies and data here, including immunofluorescence staining of both endogenous and TurboID-tagged FGF13-S (Suppl. Fig. 3), have not shown FGF13 to reside at either pre- or post-synaptic membranes. We hypothesize the synapse-related terms likely result from a role for FGF13 in trafficking of synaptic proteins to the synapse. Consistent with our hypothesis, we annotated synaptic proteins identified as FGFG13 near-neighbors as predominantly translated either in the somatic compartment, in neurites, or “other” based on a previous dataset (51) and found that 65.3% were translated in the soma compared to 14.0% that were labeled as translated in neurites (20.7% were “other”). Thirty-two “synaptic” proteins among the FGF13 near-neighbors were not present in that dataset, so 131 of 153 proteins were analyzed. When compared to the sites of translation of all synaptic proteins, as defined by the GO term “synapse” (GO:0045202) the distributions in our dataset were similar (somata: 57.3%; neuropil:14.6%; other: 28.1%) (Fig. 3B, Inset). If FGF13-S localized to the synapse the fraction of neuropil-translated proteins should be higher. Thus, most synaptic proteins identified as FGF13-S near-neighbors appear to be predominantly translated in the soma before being translocated to the synapse (Fig. 3B). This conclusion is consistent with an expanding list of roles for cytoplasmic roles for FGF13. We note that, similar to the FGF13-S dataset, 925/2025 synaptic proteins are not present in the Glock et al. dataset (51). This is likely due to the different organisms, mouse (present study) or rat (51), and different brain regions, whole brain with expression driven by NEFH promoter (present study) or CA1 region of the rat hippocampus (51).

Unexpectedly, we detected FGF13-S within LEVs and SEVs released from HEK293 cells. This was not dependent on stress, as EVs isolated from both normal and serum-deprived medium show similar results (Fig.1A). Although we attempted to remove cell debris by centrifugation to reduce contamination from FGF13-S released by dead cells, we cannot rule out that some of the detected FGF13-S in the LEV pool is due to its presence in apoptotic vesicles as the LEV pool also contains other types of large vesicles, such as large ectosomes and oncosomes. While we do not detect any FGF13 isoform in EVs from cultured mouse hippocampal neurons (Fig. 1B), its presence in EVs might depend on several factors, including cell type, tissue, stimuli and/or pathophysiological changes. Further studies are indicated to better understand the possible role of FGF13 and potentially other FHFs in LEVs. The SEV pool, which includes exosomes, ∼30-150 nm vesicles that play crucial roles in cellular communications, has been the subject of significant focus due to their tumorigenic roles (52). Further studies may clarify the presence and role of FGF13 in exosomes, but this finding is intriguing because FHFs, particularly FGF13, have been implicated in different types of cancer (53–56).

A separate study suggested that FGF12 can be secreted from cells using an FGF2-like unconventional secretion pathway in which FGF2 is first recruited to the plasma membrane via interaction with sodium-potassium ATPase (specifically subunit ATP1A1) and is then phosphorylated by TEC kinase and subsequently oligomerizes with other FGF2 molecules and undergoes self-secretion (24). Our assays would detect FGF13-S secretion from such an unconventional pathway. Further, as we detected neither the sodium-potassium ATPase nor TEC kinase in the FGF13-S TurboID dataset, we conclude that FGF13-S is not secreted via the FGF2-like mechanism.

In conclusion, this study shows that FGF13 is not secreted from HEK293 cells or mouse hippocampal neurons. Although we detected some FGF13 in extracellular vesicles from HEK293 cells by using differential centrifugation—and FGF13 in EVs could explain the reported FGF13 detected in the extracellular medium of HEK293 cells in the previous study (27)—we note that FGF13 within EVs would be incapable of binding LRR37B extracellularly. Thus, our results suggest that modulation of VGSCs and neuronal excitability depends only upon the cytoplasmic functions of intracellular FGF13-S.

## Methods

### HEK293 Cell Cultures

HEK293 human embryonic kidney cells were grown and maintained in a 5% CO_2_ incubator at 37°C in Dulbecco’s modified Eagle’s medium (DMEM; Gibco, #11995073) supplemented with 10% fetal bovine serum (FBS; VWR, #97068-085) and 1% penicillin-streptomycin (Gibco, #15140122).

### Plasmids and HEK293 cells transfection

hFGF13-S cDNA was cloned in a pcDNA3.1 background (7). The murine adipsin-HA expression plasmid (57) was kindly provided by Dr. James Lo (Weill Cornell Medicine). HEK293 cells were seeded in 6-well plates coated with poly-D-lysine (0.1 mg/mL, Gibco #A3890401) and transfected with Lipofectamine 2000 (Invitrogen, #11668019) according to the manufacturer’s instructions.

### Isolated Mouse Hippocampal Neuron Cultures

Hippocampi were dissected from P0 newborn pups and dissociated through enzymatic treatment with 0.25% trypsin (Gibco, #15090046) and subsequent trituration. For neuronal specific knockout mice, we were limited to studying hemizygous males (*Nes-cre;Fgf13^fl/Y^*) since whole body or neuronal knockout (with Nestin-Cre) is lethal before weaning (23, 58, 59), thus precluding the viability of mature hemizygous male knockouts necessary for generating female homozygous knockouts. The cells were cultured in 6-well plates for secretion experiments or plated on glass coverslips previously coated with poly-D-lysine (0.1 mg/mL, Gibco #A3890401) and laminin (20 µg/ml; Gibco, #23017015) in 24-well cell culture plates for immunofluorescence experiments. The hippocampal cells were grown in Neurobasal A medium (Gibco, #10888022) supplemented with 2% B-27 (Gibco, #17504044), 2mM L-glutamine (Gibco, #25030081) 10% fetal bovine serum (FBS; VWR, #97068-085) and 1% penicillin-streptomycin (Gibco, #15140122) in a 5% CO_2_ incubator at 37°C overnight. After 24 hours the medium was replaced by culture medium containing 2% B-27, 0.5mM glutamine, 1% FBS, 70 µm uridine (Millipore Sigma, #U3003) and 25 µm 5-fluoro-2′-deoxyuridine (Millipore Sigma, #F0503) and cultured in a 5% CO_2_ incubator at 37°C.

### Medium clearing and extracellular vesicle isolation

Twenty-four hours after transfection, HEK293 cell medium was replaced with serum-free medium (DMEM supplemented with 1% penicillin-streptomycin) or normal medium (10% serum) and cells were incubated for 24 hours in a 5% CO_2_ incubator at 37°C. The medium was then collected and subjected to sequential centrifugation steps to clear it and isolate extracellular vesicles. First, it was centrifuged for 5 minutes at 4000 x *g* on a tabletop centrifuge to remove cell debris. The supernatant was collected and centrifuged in an Optima XE-100 ultracentrifuge in a Ti rotor (Type 50.4, Beckman Coulter) for 20 minutes at 12,000 x *g* at 10°C to pellet large extracellular vesicles. The supernatant was collected and then centrifuged for 70 minutes at 100,000 x g to pellet small extracellular vesicles/exosomes. Cleared medium was collected and, for serum-deprived medium only, concentrated 30 × in an Amicon ultra centrifugal filter (10 kDa MWCO, Millipore Sigma, #UFC801024); pellets were resuspended in 100 µL of RIPA buffer (150 mM NaCl, 50 mM Tris-HCl pH 7.4, 1% Triton X-100 (v/v), 0.1% SDS (w/v), 0.5% sodium deoxycholate (w/v) 1 mM EDTA) supplemented with cOmplete protease inhibitor cocktail (Roche, # 11836170001) and 1 mM phenylmethylsulfonyl fluoride (PMSF; ThermoFisher Scientific, #36978). Pellets and cleared medium were run on a gel at the percentages of total volume indicated in Fig. 1A. Separately, cells were washed in PBS and lysed in RIPA buffer. Lysates were centrifuged at 17,000 x *g* at 4°C for 15 minutes, supernatant was collected, and protein concentration was measured by Bradford assay (Pierce, #1863028).

Isolated mouse hippocampal neurons were cultured for 12 days in 6-well plates. The medium was then collected and centrifuged at 4,000 x *g* for 5 minutes on a tabletop centrifuge to remove cell debris. Separately, cells were washed in PBS and lysed in cold RIPA buffer. Lysates were centrifuged at 17,000 x *g* at 4°C for 15 minutes, supernatant was collected, and protein concentration was measured by Bradford assay (Pierce, #1863028). Medium and lysates were run on a gel at the percentages of total volume indicated in Fig. 1B. For clusterin time-dependent secretion experiments, isolated mouse hippocampal neurons were cultured for 11 days in 6-well plates, after which the culture medium was replaced with fresh medium in half the wells and incubated for 24 hours while the other wells remained untouched. The following day medium was collected and cleared as above.

### Western Blotting

Samples were separated on Novex Tris-Glycine, 8-16% polyacrylamide gels (Invitrogen, # XP08160BOX) and transferred to PVDF membranes using the iBlot 3 Western Blot Transfer System (ThermoFisher Scientific). Membranes were blocked in blocking buffer (5% BSA (w/v), 0.1% Tween 20 (v/v)) for 1 hour at room temperature and incubated overnight with primary antibodies diluted in blocking buffer at 4°C: mouse anti-FGF13-S 1:350 (Anti-Pan-FHF-A [N235/22R]; Addgene, #190888); rabbit anti-hemagglutinin 1:1000 (anti-HA tag; Cell Signaling, #3724S); mouse anti-GAPDH 1:1000 mouse anti-Alix (Cell Signaling, #2171); rabbit anti-CD9 (Cell Signaling, #12174); rabbit anti-Syntenin (Abcam, #ab315342); rabbit anti-Calnexin (Cell Signaling, #2679); rabbit anti-FGF13 1:1000 (custom designed, YenZym); mouse anti-clusterin 1:1000 (Santa Cruz, #sc-5289). The previously described custom anti-FGF13 antibody (7) has been validated against *Fgf13* knockout models (23, 46, 48). Membranes were washed 5 times in TBST (150mM NaCl, 20mM Tris pH 7.6, 0.1% Tween 20 (v/v)) and incubated with secondary antibodies diluted in blocking buffer for 1 hour at room temperature: anti-mouse HRP 1:5000 (Santa Cruz, #sc-516102); goat anti-rabbit HRP (Cell Signaling, #7074). Membranes were washed 5 times in TBST and imaged using a ChemiDoc Touch Imaging System (Bio-Rad).

### Generation of brain specific TurboID-FGF13-S transgenic mice

hFGF13-S cDNA was fused at the 3’ of HA(x3) tag-TurboID-V5 tag cDNA to generate the TurboID-FGF13-S construct that was then cloned into the modified tetracycline-inducible promoter vector (60, 61) to generate tetracycline inducible TurboID-hFGF13-S transgenic mice in a C57BL/6 background. These mice were crossed with brain specific B6;C3-Tg(NEFH-tTA)8Vle/J (NtTA) mice (Jackson Laboratory, #025397) (62) with the human neurofilament heavy polypeptide (*NEFH*) promoter directing “Tet-Off” tetracycline-controlled transactivator protein (tTA) expression for brain-specific expression of the TurboID-FGF13-S construct. While breeding, mice were fed doxycycline (200mg/kg; BioServ #S3888) to suppress transgene expression. After weaning mice were fed normal diet to allow for transgene expression.

### *In vivo* biotinylation and sample preparation for mass spectrometry

Male and female NtTA/TurboID FGF13-S double transgenic mice, 4-8 months old, were injected daily for 3 consecutive days with Biotin (10µL/g subcutaneously, from a 2.4 mg/ml stock in PBS:DMSO 9:1) or 10% DMSO (in PBS). The mice were sacrificed to collect the brains 24 hours after the third injection. Whole brain tissues were lysed with a handheld tip homogenizer in Lysis Buffer (50 mM Tris-HCl pH 7.5, 150 mM NaCl, 10 mM EDTA, 0.5% sodium deoxycholate (m/v), 1% Triton X-100 (v/v), 0.1% SDS (w/v)) supplemented with cOmplete protease inhibitor cocktail (Roche, # 11836170001) and 1 mM phenylmethylsulfonyl fluoride (PMSF; ThermoFisher Scientific, #36978). Lysates were centrifuged at 17,000 x *g* at 4°C for 15 minutes, the supernatant was collected, and protein concentration was measured by Bradford assay (Pierce, #1863028). Biotinylation efficacy was confirmed by Western blot analysis with Streptavidin-HRP (10 µg/mL; ThermoFisher Scientific, #S911).

Brain lysates were incubated overnight at 4°C with 50 µL of streptavidin magnetic beads (Pierce, #88817), pre-washed in lysis buffer, to pull down biotinylated proteins. The next day beads were pelleted using a magnetic rack, resuspended in lysis buffer and transferred to new microcentrifuge tubes. Beads were washed 3 times with lysis buffer and 3 times with PBS. Before the final bead-pull down 10% of the resuspended beads were collected for western blot analysis and the rest submitted for mass spectrometry analysis.

### Mass spectrometry analysis

Mass spectrometric analysis was performed at the Weill Cornell Medicine Proteomics and Metabolomics Core Facility. The protein on beads was reduced with DTT, alkylated with iodoacetamide, and digested overnight with trypsin at 37°C. The digests were desalted by C18 Stage-tip columns. The digests were analyzed using a ThermoFisher Scientific EASY-nLC 1200 coupled on-line to a Fusion Lumos mass spectrometer (ThermoFisher Scientific). Buffer A (0.1% FA in water) and buffer B (0.1% FA in 80 % ACN) were used as mobile phases for gradient separation. A 75 µm x 15 cm chromatography column (ReproSil-Pur C18-AQ, 3 µm, Dr. Maisch GmbH, German) was packed in-house for peptide separation. Peptides were separated with a gradient of 5–40% buffer B over 30 min, 40%-100% B over 10 min at a flow rate of 400 nL/min. The Fusion Lumos mass spectrometer was operated in a data-independent acquisition (DIA) mode. MS1 scans were collected in the Orbitrap mass analyzer from 350-1400 m/z at 120K resolution. The instrument was set to select precursors in 45 x 14 m/z wide windows with 1 m/z overlap from 350-975 m/z for HCD fragmentation. The MS/MS scans were collected in the orbitrap at 15K resolution. Data were searched against the mouse Uniprot database (downloaded on August 7, 2021) using DIA-NN v1.8 and filtered for a 1% false discovery rate for both protein and peptide identifications. Statistical significance was determined by multiple t-tests adjusted for multiple comparisons.

### Gene Ontology (GO) analysis

GO analysis for proteins with a Log_2_(biotinylated/control)>2, P_adj_ < 0.05 was performed using g:Profiler (https://biit.cs.ut.ee/gprofiler/gost) . For each GO term, we first computed the Gene Ratio by dividing the Intersection_Size value by the Query_Size value. We then divided the Gene Ratio by (Term_Size / Effective_domain_size) to derive the Enrichment Ratio. We sorted the GO terms by the product of -log10 × P_adj_ and the Enrichment Ratio.

### Immunofluorescence

Neurons were fixed after 12 days in culture in 4% paraformaldehyde (PFA; Millipore Sigma, 1588127) for 15 minutes, washed 3 times with phosphate buffered saline (PBS) and blocked/permeabilized in 2.5% bovine serum albumin (BSA; Millipore Sigma, #A9418) with 0.2% Triton X-100 (Millipore Sigma, #T8787) in PBS for 1 hour at room temperature. Coverslips were incubated overnight with primary antibodies diluted in 2.5% BSA at 4°C: mouse anti-FGF13-S 1:250 (Anti-Pan-FHF-A [N235/22R]; Addgene, #190888); rabbit anti-Hemagglutinin 1:1000 (anti-HA tag; Cell Signaling, #3724S). Coverslips were washed 5 times in PBST (0.1% Tween-20; ThermoFisher Scientific, # J20605.AP, in PBS) and incubated with secondary antibodies diluted 1:500 in 2.5% BSA for 1 hour at room temperature: goat anti-mouse Alexa Fluor-488 (Invitrogen, #A11001); donkey anti-rabbit Alexa Fluor-568 (Invitrogen, #A10042). Coverslips were washed 5 times in PBST, incubated for 5 minutes with DAPI at room temperature, washed 3 times in PBS and mounted on glass slides (Matsunami, #SUMGP11) with mounting media (Vectashield, #H1000). Images were collected using a Leica DMi8 fluorescent microscope with Thunder imager processing.

### Data analysis

Statistical parameters are reported in the figures and figure legends. Data, graphs and images were generated with Microsoft Excel, ImageJ Fiji (63), ImageLab (BioRad), SigmaPlot, Adobe Illustrator.

### Animal studies

Mice were handled in accordance with the ethical guidelines of the National Institutes of Health Guide for the Care and Use of Laboratory Animals. This study was approved by the Weill Cornell Medical Center Institutional Animal Care and Use Committee (Protocol 2016-0042).

## Acknowledgements

We appreciate helpful comments from Manu B. Johny (Columbia University).

## Author Contributions

Designing research studies: MM, HZ, PT, DL, SOM, GSP; Conducting experiments, MM; Acquiring data, MM; Analyzing data, MM, GSP; Writing the manuscript, MM, GSP. All authors discussed the results and contributed to the final manuscript.

## SUPPLEMENTAL FIGURES

**Supplementary Figure 1:**
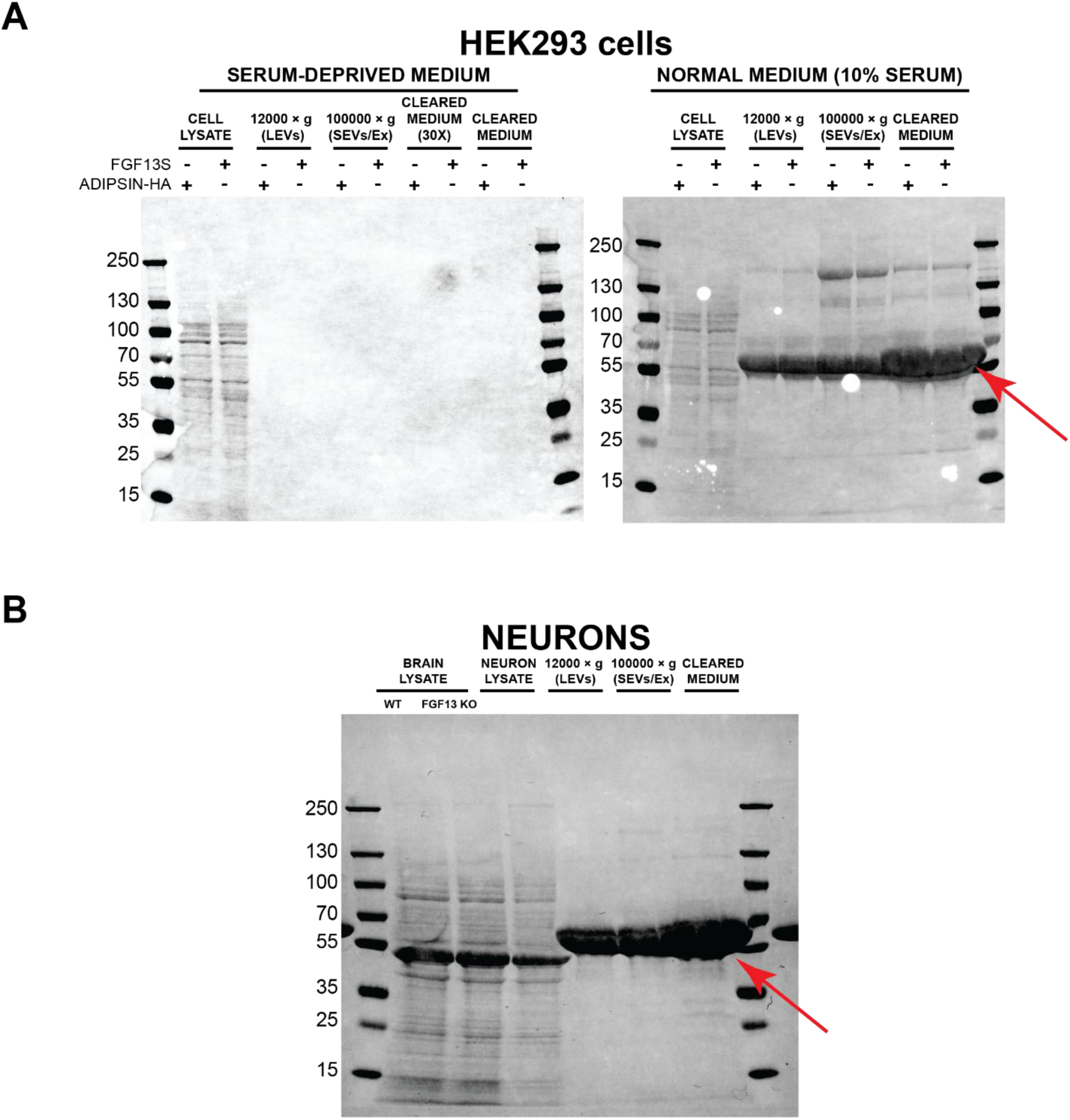
Gel artifacts caused by serum and medium supplements. Ponceau-S staining of western blot membranes showing medium clearing experiments from HEK293 (**A**) and cultured mouse hippocampal neurons (**B**) of the blots shown in Fig. 1. In the presence of 10% serum (normal HEK293 culture conditions, **A**, right) and 1% serum and 2% B27 supplement (normal neuronal culture conditions, **B**) the high amounts of albumin and other supplements cause gel artifacts (red arrows) even without medium concentration. This precludes media concentration in comparison to serum-deprived conditions (A, left) where media is concentrated 30×.

**Supplementary Figure 2.**
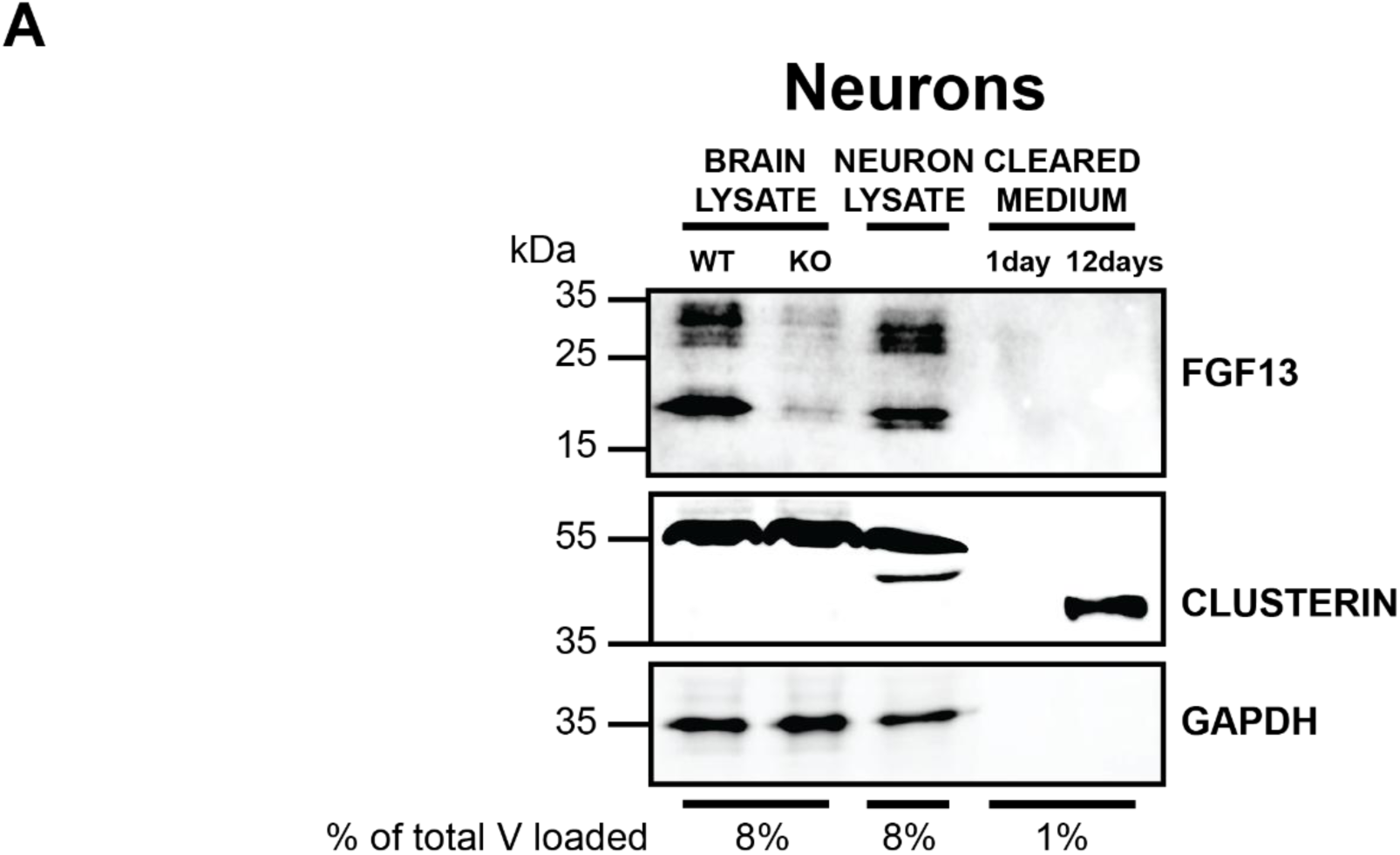
Time-dependent secretion of clusterin in cultured neurons. Western blot of cell lysates and extracellular medium derived from cultured mouse hippocampal neurons. Medium was either left untouched for 12 days or changed the day before (11th day in culture). Whole brain lysates from wild-type (WT) and *Fgf13* brain knockout (KO) mice were used to validate the detection of FGF13 isoforms. GAPDH (36 kDa) is used as intracellular marker. Clusterin is used as a positive control for secretion in neurons, and it is only detected in cells cultured for 12 days but not 1 day, consistent with its time-dependent processing and secretion. 15 µg of brain or neuronal lysate were loaded. % of total V loaded indicates the fraction of the sample loaded on the gel relative to the total volume, allowing comparison among lanes.

**Supplementary Figure 3.**
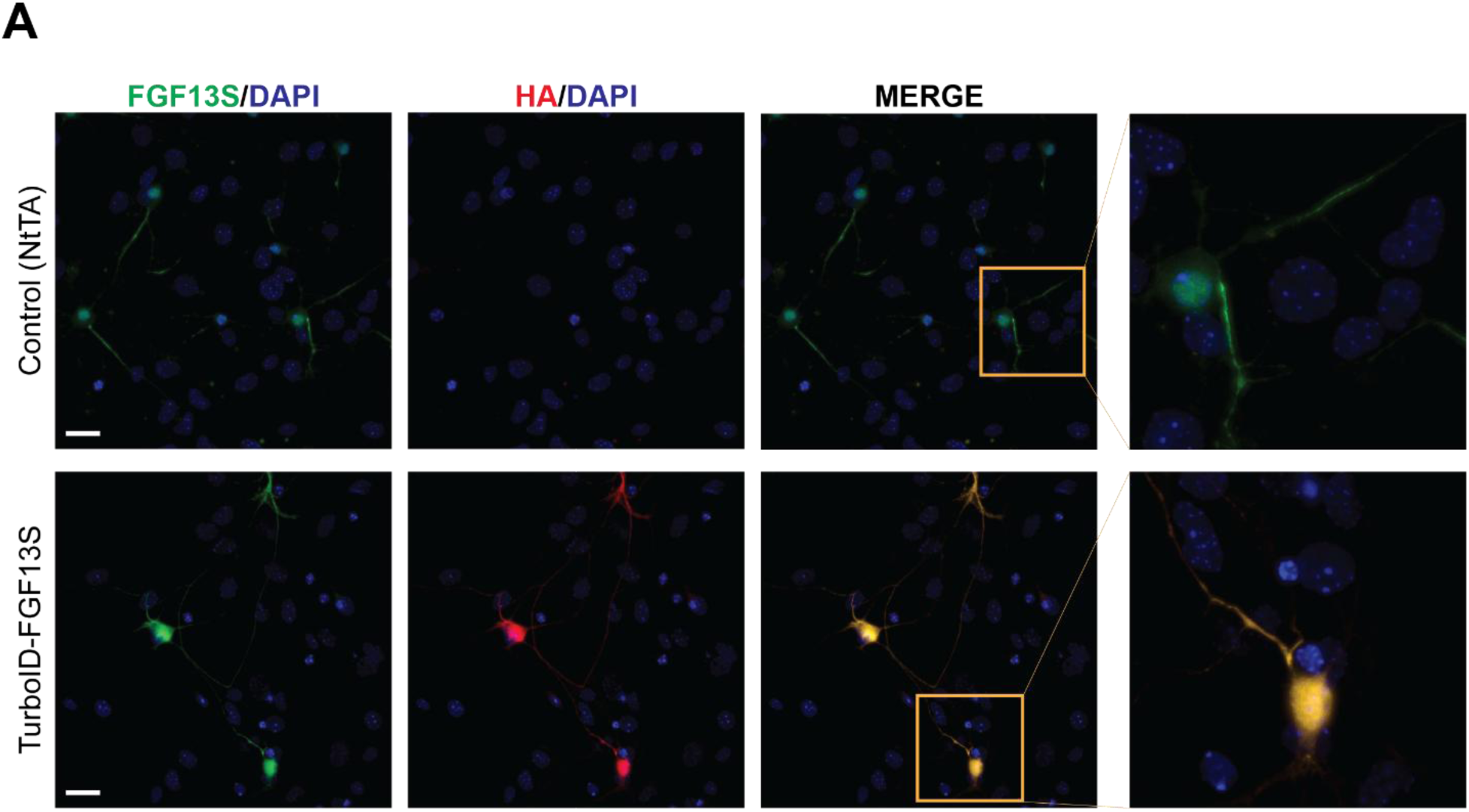
Cellular localization of the TurboID-FGF13-S transgene in mouse neurons. **A.** Representative images from immunofluorescence staining of cultured hippocampal neurons isolated from control mice (NtTA background, top) and mice expressing the TurboID-FGF13-S in the brain (bottom). Green: FGF13-S; Red: Hemagglutinin (HA) tag. Blue: DAPI. Scale bars: 30 µm.

**Supplementary Figure 4.**
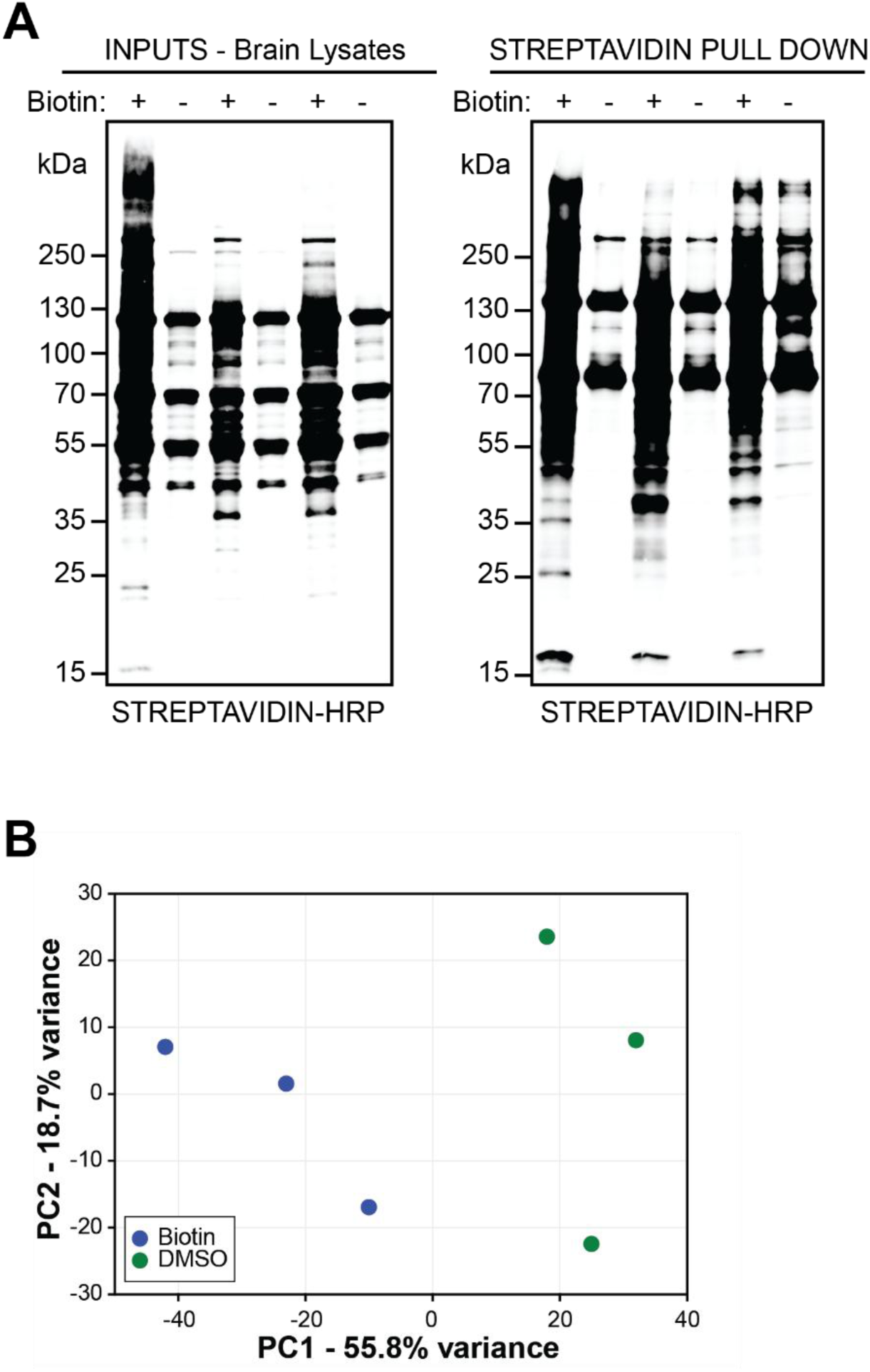
*In vivo* labeling efficiency of TurboID-FGF13S in mouse brain. **A.** Streptavidin-HRP blots of brain lysates and corresponding streptavidin pull-down samples from DMSO- or biotin-injected TurboID-FGF13-S mice, showing robust biotin labeling. The same samples were then processed and analyzed by mass spectrometry. **B.** Principal Component Analysis (PCA) plot from mass spectrometry analysis of samples shown in **A**.

**Supplementary Figure 5.**
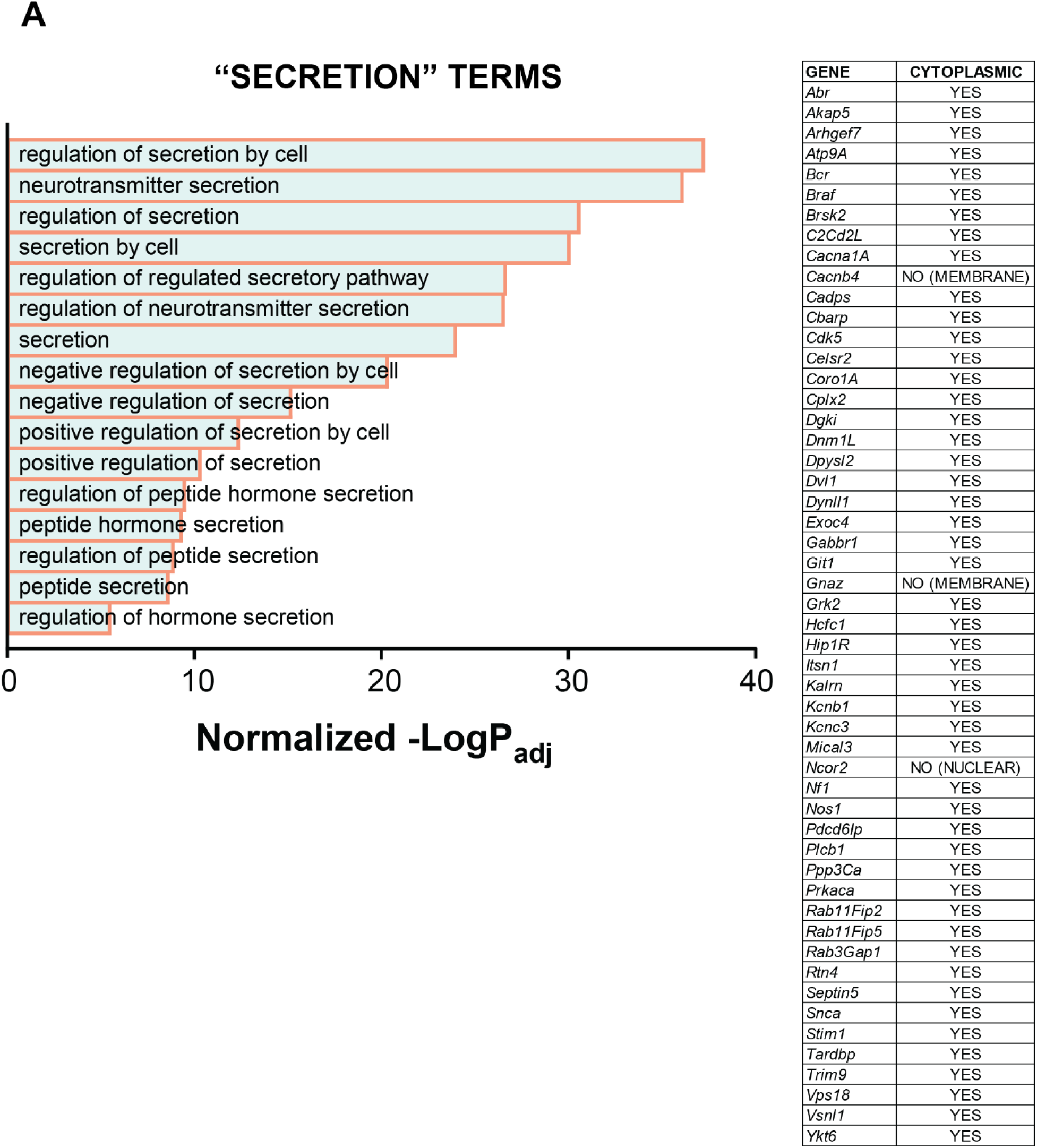
GO term analysis of “secretion” terms. **A.** The full list of GO terms with “secretion” present in their identifiers. Right table shows cytoplasmic localization of all but one protein (NCOR2). *p*-value normalization details are provided in the *Methods* section.

**Supplementary Figure 6.**
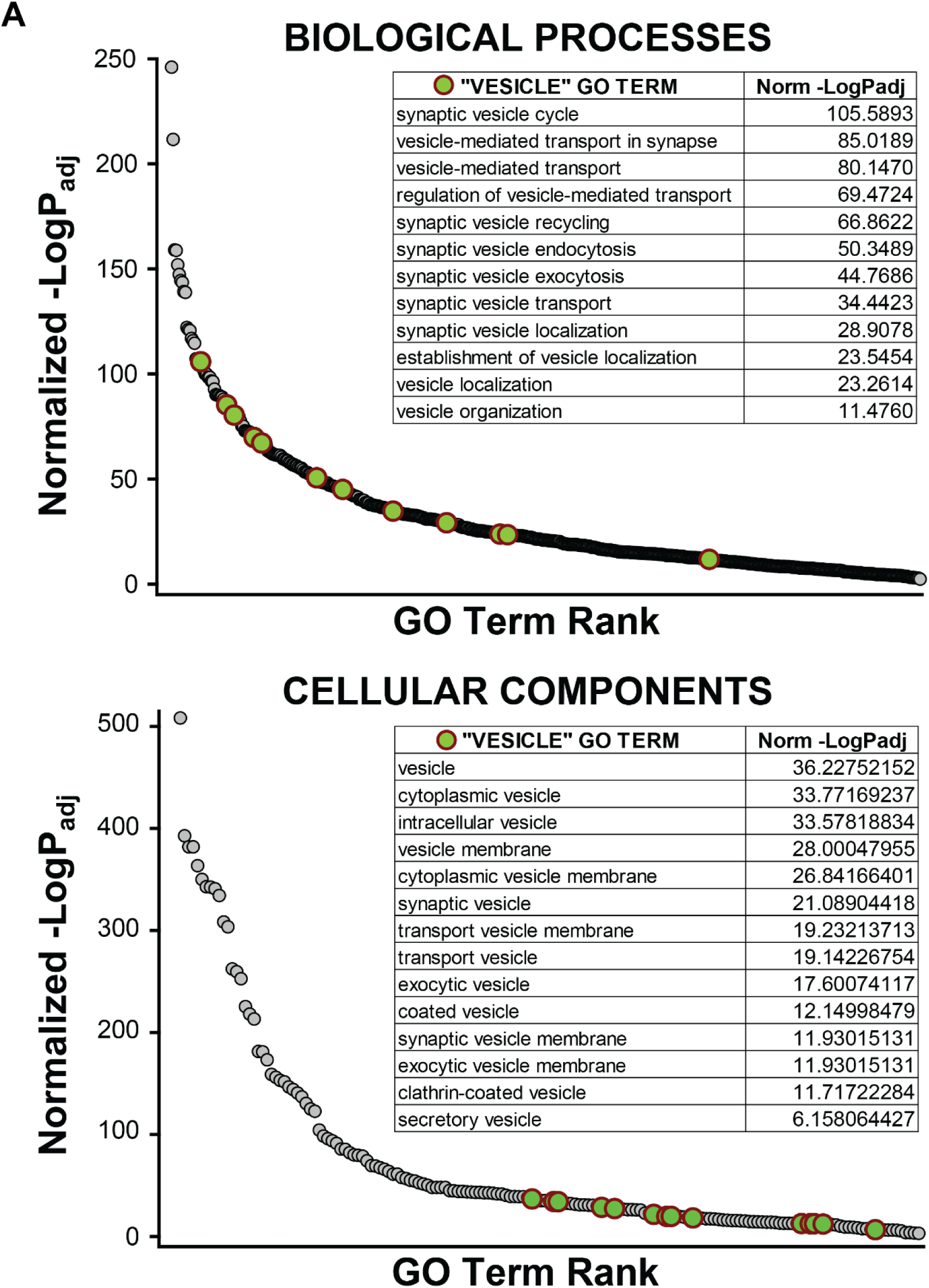
GO term analysis of “vesicle” terms. Biological Processes (**A**) and Cellular Components (**B**) GO terms (grey). Terms with “vesicles” in their identifiers are highlighted in green, and the full list is shown in the tables. The GO term rank is defined by the normalized -LogP_adj_ (descending). *p*-value normalization details are provided in the *Methods* section.

## References

1. Olsen SK, Garbi M, Zampieri N, Eliseenkova AV, Ornitz DM, Goldfarb M, et al. Fibroblast growth factor (FGF) homologous factors share structural but not functional homology with FGFs. J Biol Chem. 2003;278(36):34226–36.

2. Liu CJ, Dib-Hajj SD, Renganathan M, Cummins TR, and Waxman SG. Modulation of the cardiac sodium channel Nav1.5 by fibroblast growth factor homologous factor 1B. J Biol Chem. 2003;278(2):1029–36.

3. Wittmack EK, Rush AM, Craner MJ, Goldfarb M, Waxman SG, and Dib-Hajj SD. Fibroblast growth factor homologous factor 2B: association with Nav1.6 and selective colocalization at nodes of Ranvier of dorsal root axons. J Neurosci. 2004;24(30):6765–75.

4. Lou JY, Laezza F, Gerber BR, Xiao M, Yamada KA, Hartmann H, et al. Fibroblast growth factor 14 is an intracellular modulator of voltage-gated sodium channels. J Physiol. 2005;569(Pt 1):179–93.

5. Rush AM, Wittmack EK, Tyrrell L, Black JA, Dib-Hajj SD, and Waxman SG. Differential modulation of sodium channel Na(v)1.6 by two members of the fibroblast growth factor homologous factor 2 subfamily. Eur J Neurosci. 2006;23(10):2551–62.

6. Goldfarb M, Schoorlemmer J, Williams A, Diwakar S, Wang Q, Huang X, et al. Fibroblast growth factor homologous factors control neuronal excitability through modulation of voltage-gated sodium channels. Neuron. 2007;55(3):449–63.

7. Wang C, Hennessey JA, Kirkton RD, Wang C, Graham V, Puranam RS, et al. Fibroblast growth factor homologous factor 13 regulates Na+ channels and conduction velocity in murine hearts. Circ Res. 2011;109(7):775–82.

8. Wang C, Wang C, Hoch EG, and Pitt GS. Identification of novel interaction sites that determine specificity between fibroblast growth factor homologous factors and voltage-gated sodium channels. J Biol Chem. 2011;286(27):24253–63.

9. Yan H, Pablo JL, Wang C, and Pitt GS. FGF14 modulates resurgent sodium current in mouse cerebellar Purkinje neurons. eLife. 2014;3:e04193.

10. Pablo JL, Wang C, Presby MM, and Pitt GS. Polarized localization of voltage-gated Na+ channels is regulated by concerted FGF13 and FGF14 action. Proc Natl Acad Sci U S A. 2016;113(19):E2665–74.

11. Chakouri N, Rivas S, Roybal D, Yang L, Diaz J, Hsu A, et al. Fibroblast growth factor homologous factors serve as a molecular rheostat in tuning arrhythmogenic cardiac late sodium current. Nat Cardiovasc Res. 2022;1(5):1–13.

12. Liu C, Dib-Hajj SD, and Waxman SG. Fibroblast growth factor homologous factor 1B binds to the C terminus of the tetrodotoxin-resistant sodium channel rNav1.9a (NaN). J Biol Chem. 2001;276(22):18925–33.

13. Wang C, Chung BC, Yan H, Lee SY, and Pitt GS. Crystal structure of the ternary complex of a NaV C-terminal domain, a fibroblast growth factor homologous factor, and calmodulin. Structure. 2012;20(7):1167–76.

14. Wang C, Chung BC, Yan H, Wang HG, Lee SY, and Pitt GS. Structural analyses of Ca(2)(+)/CaM interaction with NaV channel C-termini reveal mechanisms of calcium-dependent regulation. Nature communications. 2014;5:4896.

15. Gabelli SB, Boto A, Kuhns VH, Bianchet MA, Farinelli F, Aripirala S, et al. Regulation of the NaV1.5 cytoplasmic domain by calmodulin. Nature communications. 2014;5:5126.

16. Wu QF, Yang L, Li S, Wang Q, Yuan XB, Gao X, et al. Fibroblast growth factor 13 is a microtubule-stabilizing protein regulating neuronal polarization and migration. Cell. 2012;149(7):1549–64.

17. Xiao M, Xu L, Laezza F, Yamada K, Feng S, and Ornitz DM. Impaired hippocampal synaptic transmission and plasticity in mice lacking fibroblast growth factor 14. Molecular and cellular neurosciences. 2007;34(3):366–77.

18. Bublik DR, Bursac S, Sheffer M, Orsolic I, Shalit T, Tarcic O, et al. Regulatory module involving FGF13, miR-504, and p53 regulates ribosomal biogenesis and supports cancer cell survival. Proc Natl Acad Sci U S A. 2017;114(4):E496–E505.

19. Hollern DP, Swiatnicki MR, Rennhack JP, Misek SA, Matson BC, McAuliff A, et al. E2F1 Drives Breast Cancer Metastasis by Regulating the Target Gene FGF13 and Altering Cell Migration. Scientific reports. 2019;9(1):10718.

20. Yu H, Wang H, Qie A, Wang J, Liu Y, Gu G, et al. FGF13 enhances resistance to platinum drugs by regulating hCTR1 and ATP7A via a microtubule-stabilizing effect. Cancer Sci. 2021;112(11):4655–68.

21. Munoz-Sanjuan I, Smallwood PM, and Nathans J. Isoform diversity among fibroblast growth factor homologous factors is generated by alternative promoter usage and differential splicing. J Biol Chem. 2000;275(4):2589–97.

22. Joglekar A, Prjibelski A, Mahfouz A, Collier P, Lin S, Schlusche AK, et al. A spatially resolved brain region- and cell type-specific isoform atlas of the postnatal mouse brain. Nature communications. 2021;12(1):463.

23. Lin S, Gade AR, Wang HG, Niemeyer JE, Galante A, DiStefano I, et al. Interneuron FGF13 regulates seizure susceptibility via a sodium channel-independent mechanism. eLife. 2025;13:RP98661.

24. Biadun M, Sochacka M, Kalka M, Chorazewska A, Karelus R, Krowarsch D, et al. Uncovering key steps in FGF12 cellular release reveals a common mechanism for unconventional FGF protein secretion. Cell Mol Life Sci. 2024;81(1):356.

25. Biadun M, Sochacka M, Karelus R, Baran K, Czyrek A, Otlewski J, et al. FGF homologous factors are secreted from cells to induce FGFR-mediated anti-apoptotic response. FASEB J. 2023;37(7):e23043.

26. Biadun M, Sidor S, Kalka M, Karelus R, Sochacka M, Krowarsch D, et al. Production and purification of recombinant long protein isoforms of FGF11 subfamily. J Biotechnol. 2025;403:9–16.

27. Libe-Philippot B, Lejeune A, Wierda K, Louros N, Erkol E, Vlaeminck I, et al. LRRC37B is a human modifier of voltage-gated sodium channels and axon excitability in cortical neurons. Cell. 2023;186(26):5766–83 e25.

28. Cook KS, Min HY, Johnson D, Chaplinsky RJ, Flier JS, Hunt CR, et al. Adipsin: a circulating serine protease homolog secreted by adipose tissue and sciatic nerve. Science. 1987;237(4813):402–5.

29. Welsh JA, Goberdhan DCI, O’Driscoll L, Buzas EI, Blenkiron C, Bussolati B, et al. Minimal information for studies of extracellular vesicles (MISEV2023): From basic to advanced approaches. J Extracell Vesicles. 2024;13(2):e12404.

30. Foster EM, Fernandes M, Dangla-Valls A, Hublitz P, Pangalos M, Lovestone S, et al. Glycosylated clusterin species facilitate Abeta toxicity in human neurons. Scientific reports. 2022;12(1):18639.

31. Herring SK, Moon HJ, Rawal P, Chhibber A, and Zhao L. Brain clusterin protein isoforms and mitochondrial localization. eLife. 2019;8:e48255.

32. Kolberg L, Raudvere U, Kuzmin I, Adler P, Vilo J, and Peterson H. g:Profiler-interoperable web service for functional enrichment analysis and gene identifier mapping (2023 update). Nucleic Acids Res. 2023;51(W1):W207–W12.

33. Foglio E, Puddighinu G, Fasanaro P, D’Arcangelo D, Perrone GA, Mocini D, et al. Exosomal clusterin, identified in the pericardial fluid, improves myocardial performance following MI through epicardial activation, enhanced arteriogenesis and reduced apoptosis. Int J Cardiol. 2015;197:333–47.

34. Hosseini-Beheshti E, Pham S, Adomat H, Li N, and Tomlinson Guns ES. Exosomes as biomarker enriched microvesicles: characterization of exosomal proteins derived from a panel of prostate cell lines with distinct AR phenotypes. Mol Cell Proteomics. 2012;11(10):863–85.

35. Takano T, Wallace JT, Baldwin KT, Purkey AM, Uezu A, Courtland JL, et al. Chemico-genetic discovery of astrocytic control of inhibition in vivo. Nature. 2020;588(7837):296–302.

36. Wei W, Riley NM, Yang AC, Kim JT, Terrell SM, Li VL, et al. Cell type-selective secretome profiling in vivo. Nat Chem Biol. 2021;17(3):326–34.

37. Branon TC, Bosch JA, Sanchez AD, Udeshi ND, Svinkina T, Carr SA, et al. Efficient proximity labeling in living cells and organisms with TurboID. Nat Biotechnol. 2018;36(9):880–7.

38. Smallwood PM, Munoz-Sanjuan I, Tong P, Macke JP, Hendry SH, Gilbert DJ, et al. Fibroblast growth factor (FGF) homologous factors: new members of the FGF family implicated in nervous system development. Proc Natl Acad Sci U S A. 1996;93(18):9850–7.

39. Hamdan H, Lim BC, Torii T, Joshi A, Konning M, Smith C, et al. Mapping axon initial segment structure and function by multiplexed proximity biotinylation. Nature communications. 2020;11(1):100.

40. Freal A, and Hoogenraad CC. The dynamic axon initial segment: From neuronal polarity to network homeostasis. Neuron. 2025;113(5):649–69.

41. Uhlen M, Karlsson MJ, Hober A, Svensson AS, Scheffel J, Kotol D, et al. The human secretome. Science signaling. 2019;12(609).

42. Das LT, Malvezzi M, Gade A, Matsui M, McKay M, Wei EQ, et al. FGF13 regulates cardiomyocyte impulse propagation via Cx43 trafficking independent of voltage-gated sodium channels. bioRxiv. 2025.

43. Ornitz DM, and Itoh N. The Fibroblast Growth Factor signaling pathway. Wiley Interdiscip Rev Dev Biol. 2015;4(3):215–66.

44. Schoorlemmer J, and Goldfarb M. Fibroblast growth factor homologous factors are intracellular signaling proteins. Curr Biol. 2001;11(10):793–7.

45. Hennessey JA, Wei EQ, and Pitt GS. Fibroblast growth factor homologous factors modulate cardiac calcium channels. Circ Res. 2013;113(4):381–8.

46. Wei EQ, Sinden DS, Mao L, Zhang H, Wang C, and Pitt GS. Inducible Fgf13 ablation enhances caveolae-mediated cardioprotection during cardiac pressure overload. Proc Natl Acad Sci U S A. 2017;114(20):E4010–E9.

47. Goldfarb M. Fibroblast growth factor homologous factors: canonical and non-canonical mechanisms of action. J Physiol. 2024;602(17):4097–110.

48. Gade AR, Malvezzi M, Das LT, Matsui M, Ma CJ, Mazdisnian K, et al. The NaV1.5 auxiliary subunit FGF13 modulates channels by regulating membrane cholesterol independent of channel binding. J Clin Invest. 2025;135(20).

49. Wang X, Tang H, Wei EQ, Wang Z, Yang J, Yang R, et al. Conditional knockout of Fgf13 in murine hearts increases arrhythmia susceptibility and reveals novel ion channel modulatory roles. J Mol Cell Cardiol. 2017;104:63–74.

50. Zhao R, Yan Y, Dong Y, Wang X, Li X, Qiao R, et al. FGF13 deficiency ameliorates calcium signaling abnormality in heart failure by regulating microtubule stability. Biochem Pharmacol. 2024;225:116329.

51. Glock C, Biever A, Tushev G, Nassim-Assir B, Kao A, Bartnik I, et al. The translatome of neuronal cell bodies, dendrites, and axons. Proc Natl Acad Sci U S A. 2021;118(43):e2113929118.

52. Wang G, Li J, Bojmar L, Chen H, Li Z, Tobias GC, et al. Tumour extracellular vesicles and particles induce liver metabolic dysfunction. Nature. 2023;618(7964):374–82.

53. Okada T, Murata K, Hirose R, Matsuda C, Komatsu T, Ikekita M, et al. Upregulated expression of FGF13/FHF2 mediates resistance to platinum drugs in cervical cancer cells. Scientific reports. 2013;3:2899.

54. Yu L, Toriseva M, Tuomala M, Seikkula H, Elo T, Tuomela J, et al. Increased expression of fibroblast growth factor 13 in prostate cancer is associated with shortened time to biochemical recurrence after radical prostatectomy. Int J Cancer. 2016;139(1):140–52.

55. Otani Y, Ichikawa T, Kurozumi K, Inoue S, Ishida J, Oka T, et al. Fibroblast growth factor 13 regulates glioma cell invasion and is important for bevacizumab-induced glioma invasion. Oncogene. 2018;37(6):777–86.

56. Johnstone CN, Pattison AD, Harrison PF, Powell DR, Lock P, Ernst M, et al. FGF13 promotes metastasis of triple-negative breast cancer. Int J Cancer. 2020;147(1):230–43.

57. Lo JC, Ljubicic S, Leibiger B, Kern M, Leibiger IB, Moede T, et al. Adipsin is an adipokine that improves beta cell function in diabetes. Cell. 2014;158(1):41–53.

58. Sinden DS, Holman CD, Bare CJ, Sun X, Gade AR, Cohen DE, et al. Knockout of the X-linked Fgf13 in the hypothalamic paraventricular nucleus impairs sympathetic output to brown fat and causes obesity. FASEB J. 2019;33(10):11579–94.

59. Puranam RS, He XP, Yao L, Le T, Jang W, Rehder CW, et al. Disruption of Fgf13 causes synaptic excitatory-inhibitory imbalance and genetic epilepsy and febrile seizures plus. J Neurosci. 2015;35(23):8866–81.

60. Sanbe A, Gulick J, Hanks MC, Liang Q, Osinska H, and Robbins J. Reengineering inducible cardiac-specific transgenesis with an attenuated myosin heavy chain promoter. Circ Res. 2003;92(6):609–16.

61. Hambleton M, York A, Sargent MA, Kaiser RA, Lorenz JN, Robbins J, et al. Inducible and myocyte-specific inhibition of PKCalpha enhances cardiac contractility and protects against infarction-induced heart failure. Am J Physiol Heart Circ Physiol. 2007;293(6):H3768–71.

62. Walker AK, Spiller KJ, Ge G, Zheng A, Xu Y, Zhou M, et al. Functional recovery in new mouse models of ALS/FTLD after clearance of pathological cytoplasmic TDP-43. Acta Neuropathol. 2015;130(5):643–60.

63. Schindelin J, Arganda-Carreras I, Frise E, Kaynig V, Longair M, Pietzsch T, et al. Fiji: an open-source platform for biological-image analysis. Nature methods. 2012;9(7):676–82.

